# Source to sensor coupling (SoSeC) as an effective tool to localize interacting sources from EEG and MEG data

**DOI:** 10.1101/2024.12.18.629132

**Authors:** Florian Göschl, Dionysia Kaziki, Gregor Leicht, Andreas K. Engel, Guido Nolte

## Abstract

A standard approach to estimate interacting sources from EEG or MEG data is to first calculate a coupling between all pairs of voxels on a predefined grid within the brain and then average or maximize this coupling matrix along each column or row. Depending on the chosen coupling measure and grid size this approach can be computationally very costly, in particular when a bias is supposed to be removed. We here suggest to replace this approach by a maximization of coupling between each source and the signal in sensor space. The idea is that any neuronal activity which can be estimated from recorded data must be present in sensor space in the first place. Using the imaginary part of coherency as coupling measure makes sure that we do not confuse this source to sensor coupling with a coupling of a source to itself. We found that this approach is hundreds of times faster than the conventional approach. Results for EEG resting state data indicate that the new approach has more statistical power than the conventional approach. The presentation of this specific method is augmented with a discussion of conceptual issues for various forms of vector beamformers and eLoreta.

## 1. Introduction

Neuronal oscillations have been suggested to underlie information transfer and dynamic coordination in brain networks [Singer, 1999, Engel et al., 2001, Varela et al., 2001, Fries, 2005, 2015]. These oscillations, present across a variety of frequencies, likely reflect rhythmic fluctuations in neural excitability [Buzsáki and Wang, 2012], synchronizing neuronal firing and facilitating efficient communication between distant brain areas [Fries, 2005, 2015]. The brain’s ability to dynamically form networks heavily relies on the synchronization of oscillations [Womelsdorf et al., 2007], allowing functionally specialized regions to work in concert to perform complex tasks, such as perception, attention, or memory [Siegel et al., 2012, Bowyer, 2016]. In contrast to structural connectivity, which provides insights into the anatomy of neural pathways using methods such as diffusion magnetic resonance imaging (MRI), functional connectivity describes statistical dependencies between time series of neuronal activity as measured with functional MRI, electroencephalography (EEG), or magnetoencephalography (MEG), capturing the dynamic nature of brain processes in health as well as in neurological and psychiatric diseases [Engel et al., 2013, Van Diessen et al., 2015]. Of note, there is ample evidence that combining structural and functional connectivity deepens our understanding of brain function (for reviews, see Rykhlevskaia et al., 2008, Babaeeghazvini et al., 2021).

EEG and MEG recordings feature high temporal resolution, making them suitable for the study of brain oscillations usually assessed in frequencies from around 0.1 Hz up to 150 Hz [Engel et al., 2013]. To quantify rhythmic neuronal interactions, many different metrics have been proposed in recent years, with strengths and limitations associated with each approach discussed in detail elsewhere (for reviews, see, for example, Greenblatt et al., 2012, Wang et al., 2014, Bastos and Schoffelen, 2016, Marzetti et al., 2019, Schoffelen and Gross, 2019, Basti et al., 2020).

In addition, several studies have critically assessed EEG and MEG coupling measures in sensor- and source-space using simulated as well as empirical data [e.g., Ewald et al., 2012, Haufe et al., 2013, Haufe and Ewald, 2019] and focused on caveats linked to the analysis of brain connectivity, such as the choice of forward and inverse modeling parameters [Mahjoory et al., 2017], or compared analysis pipelines allowing the aggregation of functional connectivity within physiologically defined regions of interest [Pellegrini et al., 2023].

The application of coupling methods has several pitfalls [Bastos and Schoffelen, 2016], one very prominent of them being that brain connectivity estimates are sensitive to artifacts of volume conduction/field spread. Volume conduction refers to the fact that electrical signals generated by the brain spread through the tissues of the head (with the skull having a spatial blurring effect on the electric potential distribution), causing signals from different brain sources to mix by the time they are detected by sensors on the scalp. This mixing potentially creates spurious correlations of signals not truly interacting, leading to misinterpretation of functional connectivity findings [Haufe et al., 2013, Nolte and Marzetti, 2019]. Several strategies have been suggested to mitigate the adverse effects of volume conduction, one of them projecting the complex-valued coherency (for details see Methods section below) onto the imaginary axis (imaginary part of coherency; Nolte et al., 2004), thereby explicitly removing instantaneous interactions that are potentially spurious due to field spread.

Despite the development of various methods to estimate functional connectivity that also address challenges like volume conduction [Vinck et al., 2011, Stam et al., 2007, Pascual-Marqui et al., 2011, Palva et al., 2018], there remains a need for approaches that offer both (statistical) robustness and consistency as well as computational efficiency. Traditional methods usually calculate coupling between all pairs of voxels within a pre-defined grid. Scaling with the resolution of the grid, this procedure is computationally expensive, especially when applied to large datasets. What is more, visualization and interpretation of connectivity findings between several hundreds or even thousands of sources is a difficult endeavor, let alone statistical testing in this high-dimensional space. To address these issues, we introduce a new method labeled Source to Sensor Coupling (SoSeC). This approach maximizes the coupling between each source and the signal in sensor space, eliminating the need to compute voxel-to-voxel couplings. By utilizing the imaginary part of coherency as the coupling measure, SoSeC reduces the risk of confusing source-to-source coupling with a source coupling to itself.

Our paper is structured as follows: In the*Methods* section, we introduce our new approach SoSeC, including a suggestion for bias correction, after recapitulating linear coupling measures based on the imaginary part of coherency, especially the multivariate interaction measure (MIM) and the maximized imaginary coherency (MIC) introduced by Ewald et al. [2012] and colleagues.

The *Results* section starts with findings from simulations of idealized coupled sources where we seek to illustrate differences between SoSeC, MIC, and a coupling measure based on the imaginary part of coherency that fixes source orientations such that power is maximized (which we label Fixed Direction/Univar in the following). We compare beamformer (LCMV: linearly-constrained minimum variance; Van Veen et al., 1997) with eLORETA filters (exact low-resolution electromagnetic tomography; Pascual-Marqui, 2007, Pascual-Marqui et al., 2011) as inverse methods for these and the following examples. Next, we present results on simulations concerning bias estimation and standard deviations from the coupling measures discussed here, followed by a comparison of computational costs. Before concluding, we complement our *Results* section with findings from the application of the three methods investigated to empirical (resting state) EEG data, including tests of the impact of regularization on our new method.

## 2. Methods

### 2.1. Cross-spectrum and coherence

Estimates of brain connectivity can in principle be done both in sensor and in source space. The latter requires an inverse method to estimate source signals from sensor data. For both approaches artifacts of volume conduction constitute a fundamental problem and methods are required which avoid misinterpreting incomplete demixing of empirical data as genuine brain interaction. We will use linear coupling measures based on the imaginary part of coherency [Nolte et al., 2004], which vanishes if the data, in either sensor or source space, are an instantaneous mixture of independent sources. This is exactly the case for an infinite amount of data and an approximation for a finite data set, for which significant deviations from zero must be observed to conclude that a genuine interaction was shown.

The general (though not necessary) approach to calculate coherency in sensor space is to divide data into segments or trials of a fixed length, window the data in each segment, and then calculate the Fourier transform. Let us denote the result of this *x*_*m*_(*f*) for the m.th sensor at frequency *f* with segment index omitted. The cross-spectrum is formally defined as

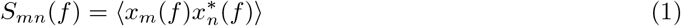

where ⟨·⟩ denotes expectation value, i.e. the hypothetical average over an infinite number of trials, which is approximated by a finite average. In the following we will omit writing the frequency dependence explicitly. Complex coherency between sensors *m* and *n* is defined as

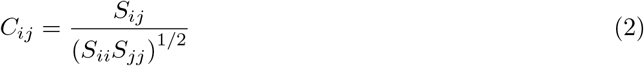

and its imaginary part is known to be robust to artifacts of volume conducting, meaning that a significant deviation from zero is inconsistent with the data being an instantaneous superposition of independent sources [Nolte et al., 2004].

### 2.2. Multivariate coupling measures

When an inverse method is applied to sensor data we (typically) get three signals for each voxel inside the brain: one for each of three orthogonal directions. In [Ewald et al., 2012] it was suggested to specify the source orientations for a given pair of voxels as the one which maximizes the ‘imaginary coherence’ (used here as a short expression for the imaginary part of coherency) between the two sources. This was formulated as a general mathematical problem which can be solved analytically in closed form. It will be recalled here as we will use the same theory albeit for a different application as will be explained below. The goal is to find two virtual signals as two linear combinations of signals of two spaces *A* and *B* with dimension *N* and *M*, respectively.

We write the corresponding data in the frequency domain as vectors *x*_*A*_ = (*x*_1_ … *x*_*N*_)^*T*^ and *x*_*B*_ = (*x*_*N*+1_ … *x*_*N*+*M*_)^*T*^. In the case of three dimensional dipole moments at given brain voxels we have *N* = *M* = 3.

The cross-spectrum between all components of 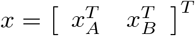 exhibits the block form

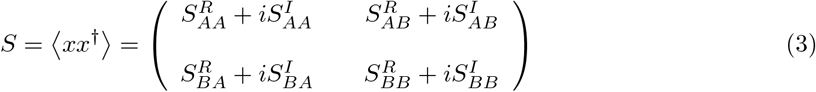

where † denotes the conjugate transpose and *S*^*R*^ denotes the real and *S*^*I*^ the imaginary part, respectively.

The goal is now to construct signals as linear superpositions of the two subspaces, namely

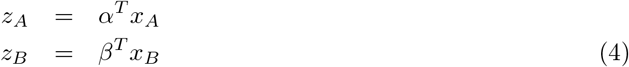

with two column weight vectors *α* and *β* of size *N* and *M*, respectively, such that the imaginary part of coherency between *z*_*A*_ and *z*_*B*_ is maximized. This quantity is given by

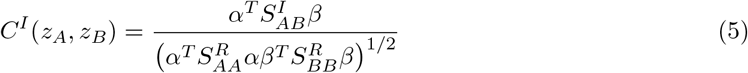

Maximization of the right hand side of (5) can be done analytically. We here only present the solution, and refer to [Ewald et al., 2012] for the derivation.

We first construct a matrix

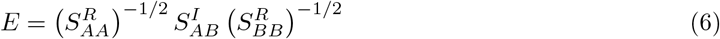

and then solve the eigenvalue equation

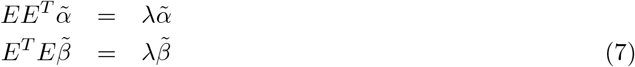

Note, that the eigenvalues, but not the eigenvectors, are identical for *EE*^*T*^ and *E*^*T*^ *E*. Let’s call *λ*_*max*_ the largest eigenvalue with eigenvectors 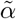 and 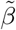,and then

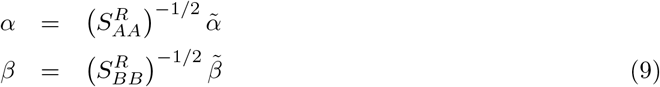

and the maximizing weight vectors are related to the eigenvectors as

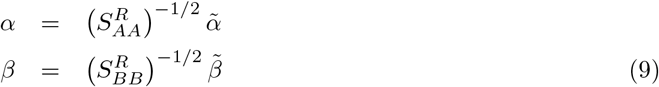

In this work we only need the eigenvalues and not the eigenvectors. Then we only need to solve one of the eigenvalue equations, of course preferably the one with smaller dimension.

### 2.3. Source to sensor coupling (SoSeC)

A natural application is the estimation of coupling between two voxels where each voxel contains signals in three dipole directions. In that case the subspaces *A* and *B* are both 3-dimensional. If we calculate coupling between all pairs of voxels for *P* voxels, we need to solve *P*^2^ eigenvalue equations which is computationally very costly. The new point here is the application and not the theory itself. We propose to take as subspaces the 3-dimensional subspace for a given voxel and the M-dimensional sensor space. We refer to this as SoSeC, the source to sensor coupling. The rationale behind this is that any signal which can be recovered on source level from sensor data must have been in the sensors in the first place. The maximal coupling between a voxel and sensors indicates whether that voxel interacts with something else, regardless of what the other source is. Most of all, the computational cost is much lower, because we only need to solve *P* eigenvalue equations rather that *P*^2^. Furthermore, if we study pairs of voxels, we implicitly assume that both sources are dipolar, whereas with SoSeC only one of the sources, the voxel, is assumed to be dipolar. We will illustrate this point in simulations below. We note, that this approach would have been meaningless if we would have taken coherence, rather than imaginary coherence, as coupling measure, because the signal originating in each voxel is also contained in the sensor signal and we would always observe the coupling of a signal with itself. Maximal coherence is always equal to 1 in this case.

In the final part of this section we will present equations on how to calculate all relevant crossspectral matrices from the cross-spectrum at sensor level and from the spatial filters used for inverse calculations. Let *L* be a 3 × *M* matrix corresponding to the spatial filter used to estimate the source activity in a specific voxel for 3 dipole directions from the signal in *M* sensors. This means, that if *x*_*A*_ are the data in sensor space in the Fourier domain for a given segment, then

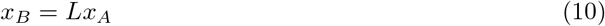

are the corresponding data at source level for that specific voxel. Standard methods to construct such filters are LCMV-beamformers or eLoreta, which will be both used in this work, but the theory is applicable to any inverse method as long as the mapping from sensors to sources is linear. Furthermore, let *S* be the *M* × *M* cross-spectral matrix at sensor level. Then the calculation of the matrices in (3) for SoSeC is straight forward

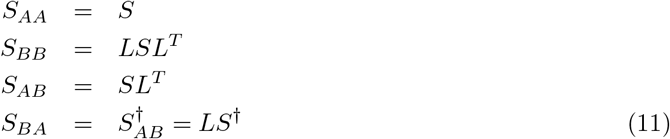

For the calculation of MIC (see previous section) we need two spatial filters for two voxels denoted as *L*_1_ and *L*_2_ and the corresponding matrices in (3) read

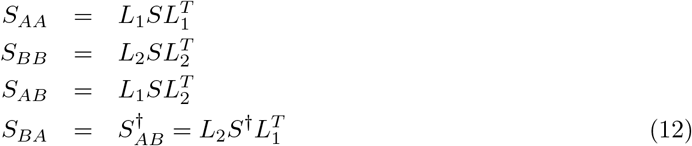

Both SoSeC and MIC are multivariate methods. Below we also compare results with dipole direction fixed by maximizing the power for each voxel. Then imaginary coherence for a given voxel pair is calculated for two univariate signals, one for each voxel, which does not require the optimization of coupling with respect to the dipole orientation. In the following, we refer to this case as ‘univar’ or ‘fixed dir’.

### 2.4. Bias and standard deviation

Both for SoSeC and for MIC we maximize with respect to weight vectors, which introduces a bias in the coupling measures towards larger values. This bias is larger for SoSeC compared to MIC because we maximize over more parameters. A small bias is also contained for the imaginary part of coherency because in the end we take the absolute value.

We here use the standard Jackknife procedure to estimate bias and standard deviation on a single trial level. Let us first recall the basis principles. Let *z* be some quantity estimated from *N* independent observations *x*_1_, …, *x*_*N*_ with some function *f*

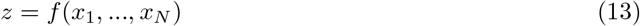

If *z* is biased for a finite number of observations this bias is larger if we have fewer observations, in particular if we take out just one of the observations. Taking out the *i.th* observation we get new estimates in the form

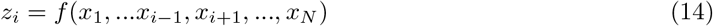

With a bias, the average of *z*_*i*_ is systematically different from *z* and using this, a debiased estimate can be constructed as

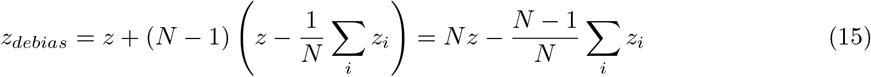

The quantities *z*_*i*_ can also be used to calculate an estimated standard deviation *σ* both of *z* and *z*_*debias*_ as

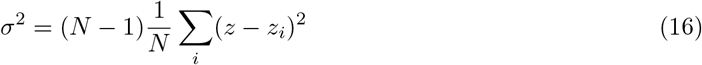

We emphasize that for such estimates it is assumed that the observations are independent. This is not the case e.g. if we consider consecutive segments of 1 second duration in EEG resting state data. Neighboring segments typically have a substantial correlation. As an alternative we suggest to take longer segments of about 10 seconds duration such that neighboring segments are at least approximately independent. This will be tested below in simulations.

The Jackknife procedure is based on a Taylor expansion around the true value up to quadratic terms and it is assumed that the estimates are close to the true values. It is important that the Taylor expansion up to second order is a good approximation in the vicinity of the true value. To illustrate this, an approximation can be poor if e.g. the estimate is an absolute value of an average, because the absolute value is not a differentiable function at the origin, whereas it is perfect if the square of the average is taken. We found a similar behavior here: we can successfully correct for the bias if we use the square of imaginary coherence as coupling measure but not if we use imaginary coherence itself. This will be shown in simulations below.

### 2.5. General remarks on forward and inverse calculations

#### 2.5.1. Forward calculation and Toolbox

For the forward calculation we assumed a standard MNI head model with a 3-shell realistic volume conductor. The forward problem is solved with a semi-analytic expansion of the EEG lead field as proposed in [Nolte and Dassios, 2005]. The head model and the code for all forward and inverse methods used here are contained in the ‘MEG and EEG toolbox of Hamburg’ (METH) which can be download at https://www.uke.de/english/departments-institutes/institutes/neurophysiology-and-pathophysiology/research/research-groups/index.html.

#### 2.5.2. Beamformers

A beamformer is a spatial filter which maps activity in sensor space to estimated activity in source space. We here use a formulation essentially identical to the vector beamformer proposed in [Van Veen et al., 1997] and usually called LCMV (linearly constrained minimum variance) beamformer. We refer to this beamformer as the standard beamformer.

Let *L* be the *M* × 3 matrix for *M* sensors and 3 source directions denoting the forward mapping for some voxel in the brain, i.e. *L*_*mk*_ is the electric potential in sensor *m* of a unit dipole located in that voxel in direction *k*. Further, let *C* = *C*_*R*_ + *iC*_*I*_ be the cross-spectrum in sensor space with *C*_*R*_ and *C*_*I*_ being its real and imaginary parts, respectively. We now construct a real valued spatial filter *A*, which is an *M* × 3 matrix and estimates data in source space *y* from data in sensor space *x* as *y* = *A*^*T*^ *x*. With *Tr* denoting trace it is designed to minimize the total power in that voxel

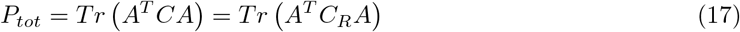

subject to the constraint

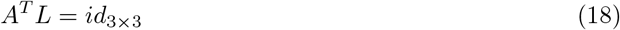

where *id*_3*×*3_ is the 3 × 3 identity matrix. This choice maximizes the signal to noise ratio if the signal in that voxel is independent of all other activities. It is well known, that beamformers can perform poorly if that assumption is substantially violated.

The minimization problem can be solved analytically as

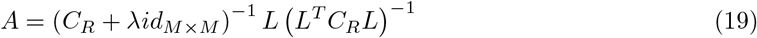

where we have included a fixed regularization with *λ* = 0.05*Tr*(*C*_*R*_)*/M*.

The real part of the cross-spectrum is identical to the covariance matrix of the filtered data apart from possible deviations due to different filter settings. The proof of this is rather lengthy and omitted here. It is analogous to similar proofs in [Bruns, 2004, Nolte et al., 2020]. Therefore, apart from such conceptually meaningless deviations, the beamformer in (19) is equivalent to the standard LCMV beamformer.

A slightly different approach was proposed in [Gross et al., 2001] where *C*_*r*_ in (19) is replaced by the full cross-spectrum *C*. The corresponding beamformer is usually called ‘DICS-beamformer’. A drawback is that the filter itself is in general complex. Although one can just set the imaginary part of the filter to zero, this is not equivalent to starting with the real part of the cross-spectrum in the first place which follows from minimizing *P* and only allowing real valued filters.

Let us now discuss the linear constraint in (18). Although to our knowledge it is never called as such, such a beamformer is really a ‘nulling-beamformer’. In nulling beamformers additional constraints are set to remove contributions from specified other regions in the brain entirely [Hui et al., 2010]. What is removed in the standard LCMV beamformer are not contributions from other regions but from orthogonal directions. This is not necessary if the goal would be just to estimate the source activity as a vector, i.e. for all three orthogonal source directions. One could also construct filters for each dipole direction separately by replacing in (19) *L* by the k.th column of *L* for *k* = 1, 2, 3 resulting in filters for each dipole component without nulling out orthogonal directions. However, a severe problem arises with such a beamformer because results would then depend on the orientation of the coordinate system. If we express a beamformer in a generic form as *A* = *f* (*L*) then results are independent of the orientation of the coordinate system if for any orthogonal 3 × 3 matrix *O* we have

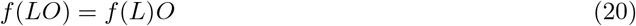

This is the case for the standard LCMV-beamformer, which follows directly by replacing *L* with *LO* in (19), but not for a vector beamformer without nulling out orthogonal directions, which can be checked numerically by the reader with a few examples. Hence, the rotational invariance justifies the standard vector beamformer, which was, to our knowledege, not mentioned before.

#### 2.5.3. eLoreta

While the spatial filters constructed as beamformers depend both on the forward model and on the data via the cross-spectrum, eLoreta is a filter which only depends on the forward model. Its construction is rather complicated and we refer to the original papers [Pascual-Marqui, 2007, Pascual-Marqui et al., 2011] for details. The filter is found from an iteration which may take a few seconds until convergence with details depending on the number of channels, the number of voxels and the stopping criterion. However, since it does not depend on the data, it has to be calculated only once for a given forward model. The filter has the remarkable properties that a) the full inverse solution explains the data, and b) for single dipoles the maximimum of the estimated source density coincides with the true location of that dipole. In contrast to beamformers the eLoreta-filter cannot be calculated separately for each voxel. We tested rotational invariance numerically using random numerical examples. We found that relative deviations are in the order of 10^−6^ which corresponded to the stopping criterion of the iteration. In general, eLoreta solutions are more blurred than beamformer solution but we found them also to be statistically more robust as we show below. Also, eLoreta does not have the conceptual problem of beamformers in this context, namely that we study brain interactions while beamformers are constructed assuming that the reconstructed activity in each voxel is independent of any other activity.

## 3. Results

### 3.1. Simulations

#### 3.1.1. Illustrative example

To illustrate the conceptual differences between the various coupling methods we simulated three idealized examples. In first example we placed two sources pointing in y-direction at coordinates *x* = ± 4*cm, y* = −2.5*cm* and *z* = 5*cm* in an MNI coordinate system. They are shown in the top left panel of Fig.1 as blue arrows. The time course of the left source was chosen as white noise filtered at 10Hz with a bandwidth of 1Hz. The time course of the right source was identical to the one of the left source apart from a delay of 0.012 seconds corresponding to a phase difference of about 45 degrees. Thus, the absolute value of coherence between the sources was 1, and its imaginary part approximately 0.7. To the corresponding signal in sensor space, we added noise generated from independent sources equally distributed in the entire brain and also filtered at 10Hz. The signal to noise ratio, defined as the ratio of the Frobenius norms of signal and noise, respectively, was chosen to be 1. As inverse method we chose the beamformer. Results for eLoreta are qualitatively similar but more blurred. (Below we will also show results for both beamformer and eLoreta for empirical data.)

**Figure 1:**
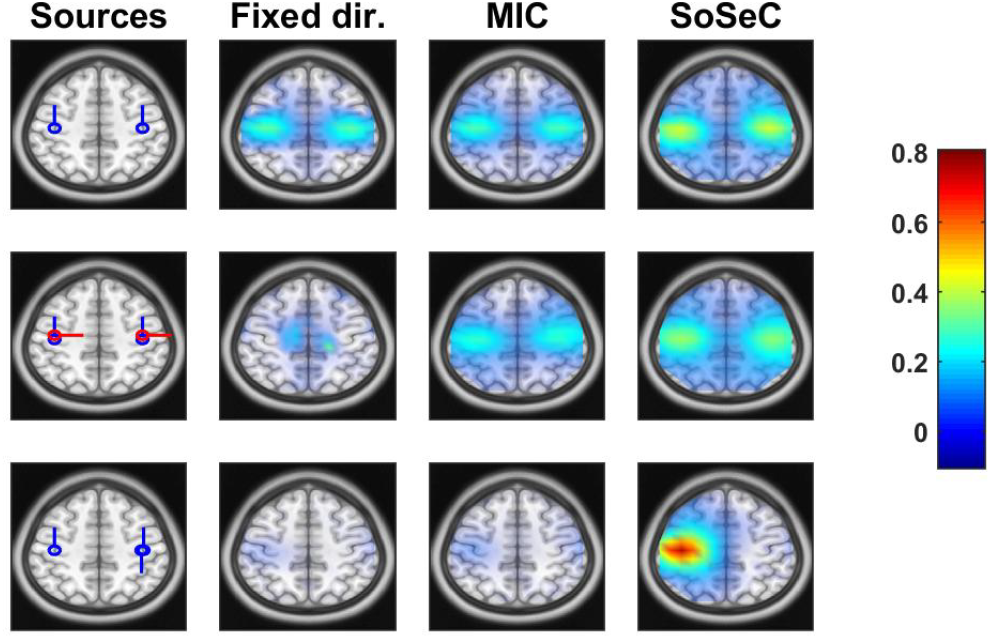
Simulation results for three different coupling measures and three different source scenarios. Upper row: two coupled dipolar sources plus additional noise. Middle row: two coupled dipolar sources plus two uncoupled sourced at the same location plus additional noise. Lower row: Coupled dipolar and quadrupolar source plus additional noise. Left column: illustration of coupled (blue) and uncoupled (red) sources. Columns 2-4: coupling measures for three different methods. Note that for the second and third column the coupling is a matrix for all voxel pairs and we show for each column the maximum across rows.

Results for this simulated scenario are shown in the upper row of Fig.1. In the second column we show results for fixed dipole directions with orientations fixed by maximizing power. We have first calculated imaginary coherence for each pair of voxels in source space and then maximized for each voxel over all other voxels. In the third column we use the multivariate method MIC for each pair, i.e. dipole directions are chosen to maximize imaginary coherence for each pair separately and then we again maximize for each voxel the coupling over all second voxels. Finally, in the fourth column we show results for the proposed method SoSeC. For all results we removed the bias as explained in section 2.4. We observe comparable results for all three coupling methods.

In the second example we added additional uncoupled sources at the same origins as the coupled sources but pointing to x-direction and again with filtered white source as signal amplitudes but now independent of each other. These additional sources are shown as red arrows in the middle panel of the left column. In sensor space, the Frobenius norm of these additional sources was twice as large as the the one of the original two sources. Additional noise was added as in the first example with signal to noise ratio referring to all four dipoles. Results are shown in the second row. We observe that the method with fixed dipole direction fails for this example. Source directions are dominated by the non-interacting sources and miss the coupling of the weaker coupled sources.

In the third example we replace the right source dipole by a source quadrupole as shown by the blue arrows in the left panel of the third row while everything else was identical to the first example. We observe that both fixed direction and MIC fails completely, while SoSeC is able to detect the left dipolar source but not the quadrupolar source on the right. In contrast to MIC, where both sources have to be dipolar to lead to observable coupling, for SoSeC only one of the sources has to be dipolar.

#### 3.1.2. Bias and standard deviation

We tested formulas given in (15) and (16) for the estimation of bias and standard deviation for the coupling measures discussed here. In the jackknife procedure small portions of the data are excluded. Equations are based on the assumption that these portions are statistically independent of the rest of the data. This is a valid assumption in an event related experimental design where each portion corresponds to a single trial. However, in resting state data, which are analyzed here, one must be careful not to choose portions too small such that temporal correlations are larger than the portion size that leads to systematic misestimations. We simulated the same situation as reported in the first example above, i.e. two coupled dipoles containing oscillations at 10 Hz plus additional noise. The portion duration was 5 seconds. We show results for the beamformer. Results for eLoreta were similar.

The jackknife procedure is based on Taylor expansions of the corresponding coupling measure around the true value, whereas ‘true value’ here means the result for a hypothetical infinite amount of data. For this procedure it is assumed that the total amount of data is sufficiently large such that the estimate is close to this true value such that a first order Taylor expansion is a reasonable approximation. It is important to note that the jackknife procedure becomes inaccurate if the analyzed quantity is not differentiable at or near the true value. This is the case if we analyze e.g. the absolute of the imaginary part of coherency rather than the square of it. We will therefore analyze coupling measures both as absolute values and as the squares of them, and will show that we only get sufficiently accurate results when analyzing the squares.

We simulated a single data set consisting of 5000 minutes of data and considered results for this data set as the true results. In addition, we simulated 100 data sets each consisting of 5 minutes. For each of these data sets and for each voxel we calculated the coupling measures with and without bias removal and estimated the variance with the jackknife procedure. We finally averaged these results over the 100 data sets. In Fig.2 we show results for the three different coupling measures (univariate, MIC, SoSeC) using unsquared and squared versions of the coupling measures. We show the average estimates with and without bias removal as a function of the true coupling. We observe that the bias is accurately removed only if we use the squared version of the coupling measure. The bias itself is negligible only for the univariate case and largest for SoSeC in particular for small coupling.

**Figure 2:**
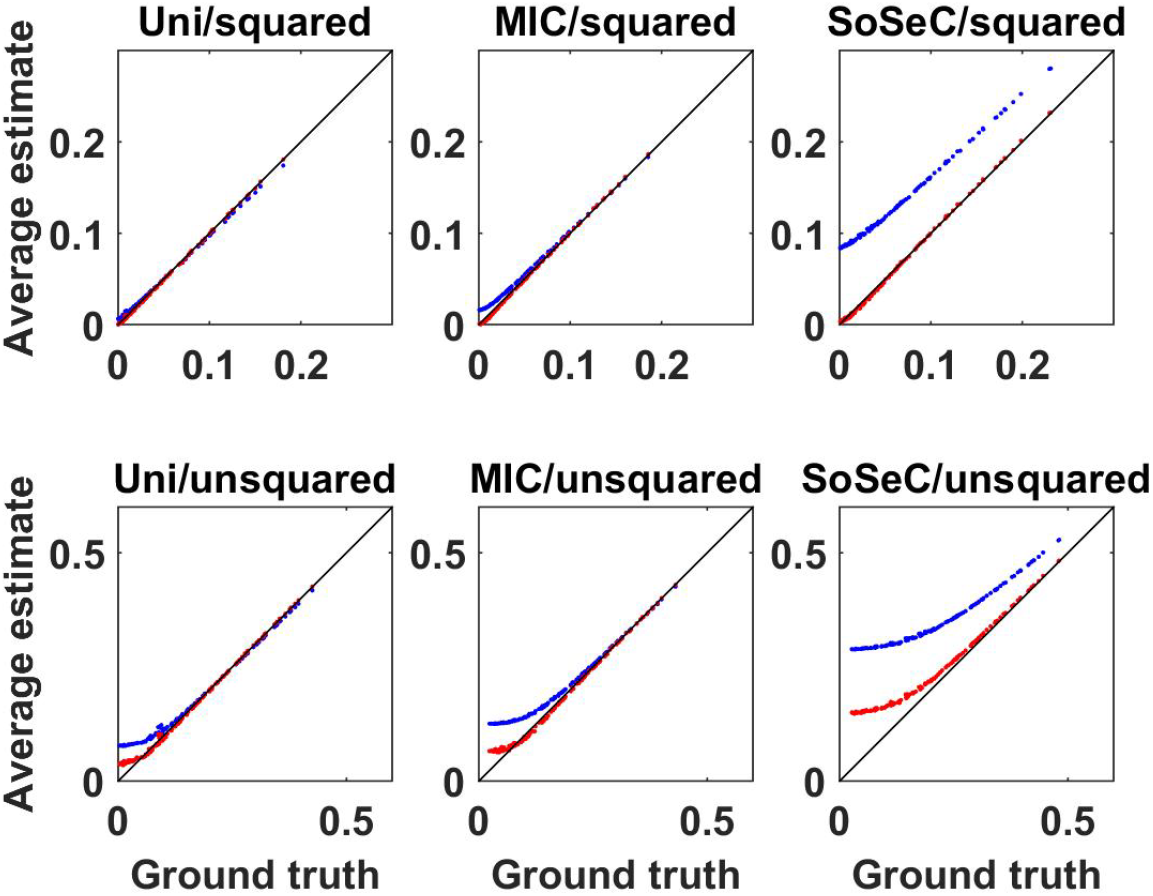
Mean of original (blue dots) and bias removed (red dots) coupling for different coupling measures as indicated as functions of ‘true’ results, i.e. results for a single extremely large data set.

In Fig.3 we show the average of the estimated standard deviation for the 100 data sets as a function of the standard deviation found directly from the 100 results. We observe that these estimates are less accurate than the bias estimates but work reasonably well in particular for SoSeC using the squared coupling measure.

**Figure 3:**
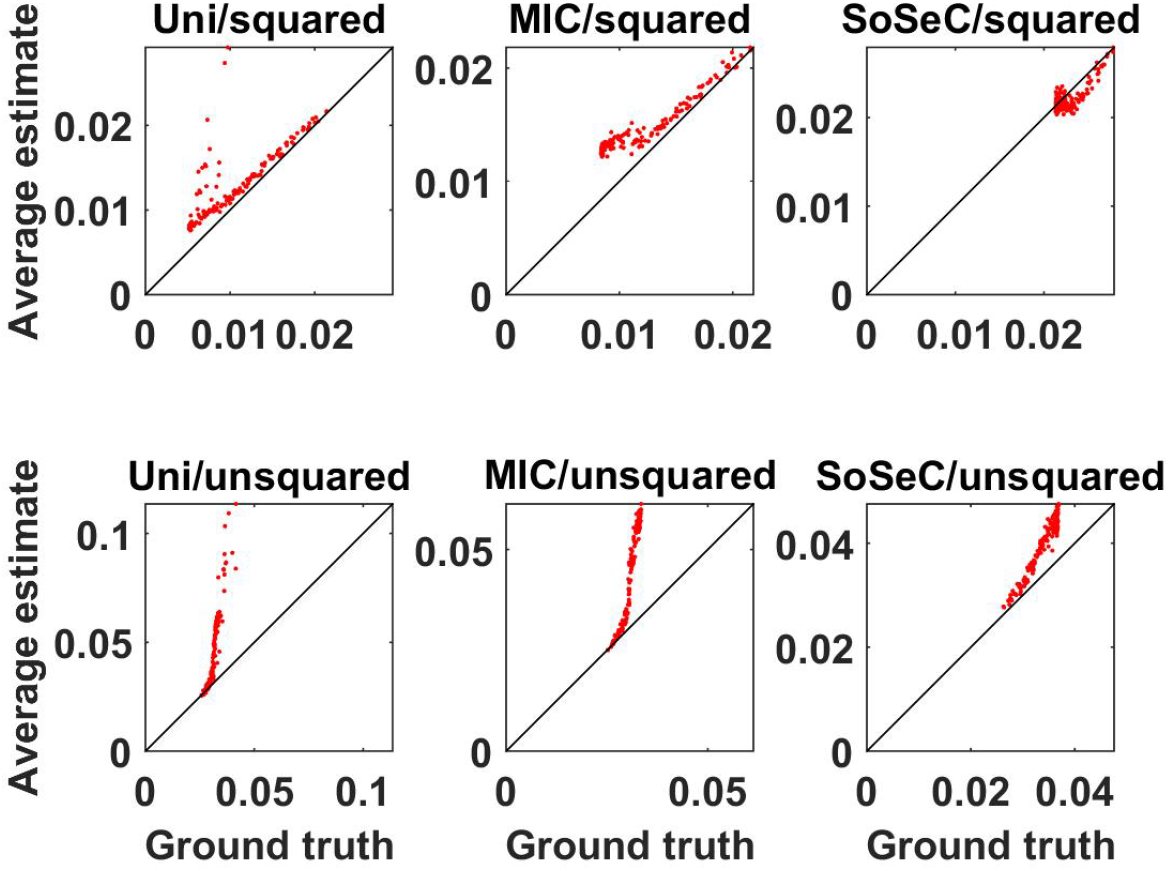
Mean of estimated standard deviations for different coupling measures as indicated as a function of ‘true’ standard deviation, i.e. standard deviation calculated across all results.

#### 3.1.3. Computational cost

For *N* voxels in the brain, the proposed method requires only *N* optimizations, while MIC requires *N*^2^ optimizations leading to substantial computational advantage of SoSeC compared to MIC. This is the case even though for SoSeC each optimization, which can be done analytically, is computationally more involved as there are *M* + 3 unknown parameters for *M* sensors while there are only 6 unknown parameters for MIC. However, the relative advantage of SoSeC over MIC still depends both on the numbers of sensors and on the number of voxels. In Fig.4 we show the computational costs (in seconds of computation time) for an MEG system with 271 channels and for an EEG system with 61 channels as a function of the number of voxels for all three methods (SoSeC, MIC, and fixed direction) compared here. For this calculation we used a beamformer. We observe that for more than 1000 voxels SoSeC is several hundred times faster than MIC and even for as few as 100 voxels SoSeC is still around 10 times faster than MIC. On the other hand, calculations with fixed direction are much faster than both MIC and SoSeC because here we do not need any optimization at all and the coupling matrix for all pairs of voxels can be calculated in a single analytical step. If computational cost is the only criterion for the choice of method the fixed direction is the best by far. However, we will show in illustrative examples of empirical data that results for multivariate methods are much more convincing.

**Figure 4:**
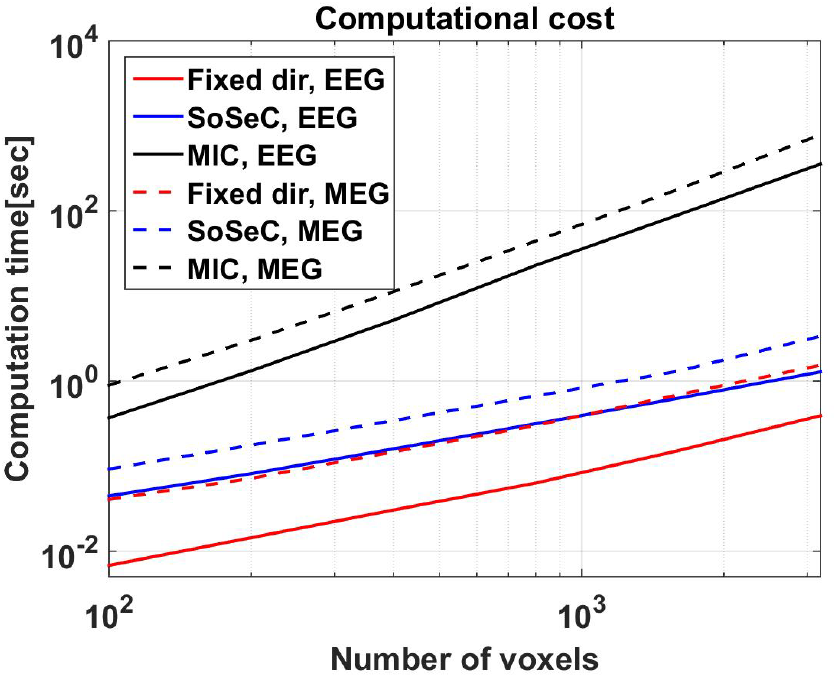
Computational cost for the three coupling measures (fixed direction, MIC and SoSeC) as function of the number of voxels for an EEG system with 61 channels and an MEG system of 271 channels with a beamformer as inverse method. Using a fixed direction is much faster than the multivariate methods MIC and SoSeC but may result in poor estimations as will be shown below. Depending on the number of voxels SoSeC can be hundreds of times faster than MIC.

### 3.2. Empirical data

#### 3.2.1. An example data set for EEG resting state data

For illustrative purposes, we first present results for a single data freely available from the METH toolbox. The data set consists of 5 minutes resting state activity measured from a healthy participant with 64 EEG channels under eyes closed condition. It is one data set out of 24 sets (of healthy control participants plus another 22 data sets of patients recorded at the University Medical Center Hamburg-Eppendorf) which are described below in more detail.

We first calculated the imaginary part of coherency on sensor level using segments of 1 second duration, corresponding to a frequency resolution of 1Hz, with 50% overlap. Results for all pairs of channels as a function of frequency are shown in Fig.5. Vertical lines were set manually indicating clear peaks of the imaginary part of coherency. The corresponding frequencies are reasonable candidates for further analysis of brain coupling on source level.

**Figure 5:**
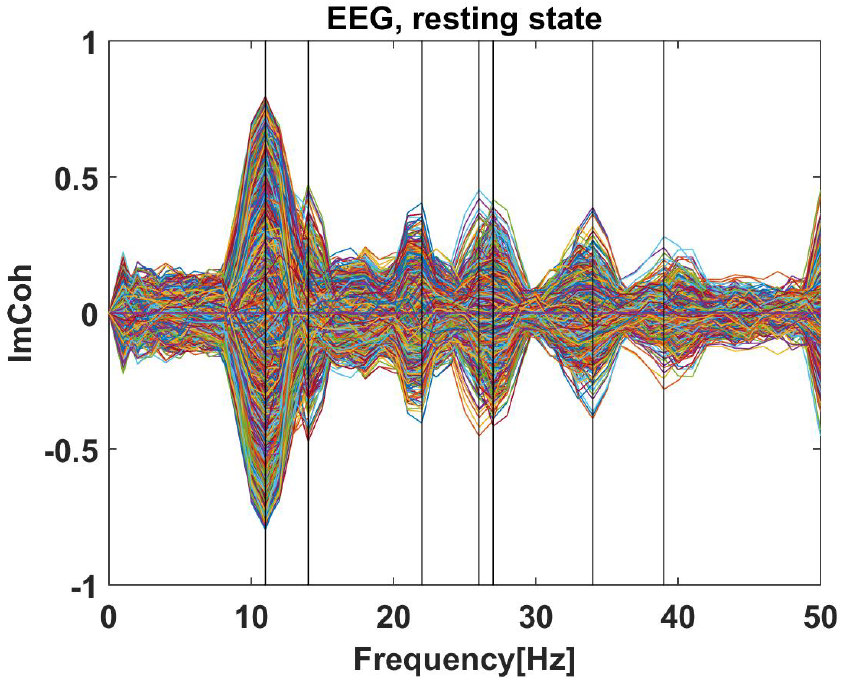
Imaginary coherence as a function of frequency for all channel pairs for one example EEG resting state data set. Vertical lines denote peaks and correspond to candidate frequencies for further analysis of brain interactions.

We further analyzed the data using 2113 voxels equally distributed in the entire brain with 1cm between neighboring grid points. For a more compact visualization we extrapolated results on the cortical surface using a Gaussian kernel with width *d* = 5*mm*. Specifically, let *r*_*i*_ be the location of the *i.th* grid point, *r*_*j*_ be the location of the *j.th* cortical point, and *c*_*i*_ the coupling measure for the *i.th* grid point, then the coupling at the *j.th* cortical point is calculated as a weighted sum

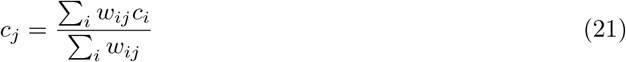

with

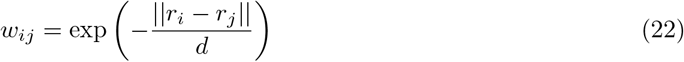

Calculations and visualizations were done in an MNI template head using a semi-analytic method for EEG forward calculation in a three shell volume conductor of realistic shape Nolte and Dassios [2005].

In Fig.6 we show results for all three coupling measures considered here, using both the beamformer and eLoreta as inverse method at 11 Hz. The bias was removed in all cases. We observe that results for MIC and SoSeC are very similar while results for fixed direction are substantially smaller and show a less clear topographic distribution. More important than the coupling values themselves are the corresponding z-scores, defined by the coupling value divided by the estimate of the standard deviation, which represent the stability of the coupling values and which are shown in Fig.7. We observe substantially larger z-scores for SoSeC than for MIC and substantially larger z-scores for MIC than for fixed direction. Also, eLoreta leads to higher z-scores than a beamformer. As a final example we show z-scores for the signal at 14Hz in Fig.8, again with highest values for SoSeC in combination with eLoreta.

**Figure 6:**
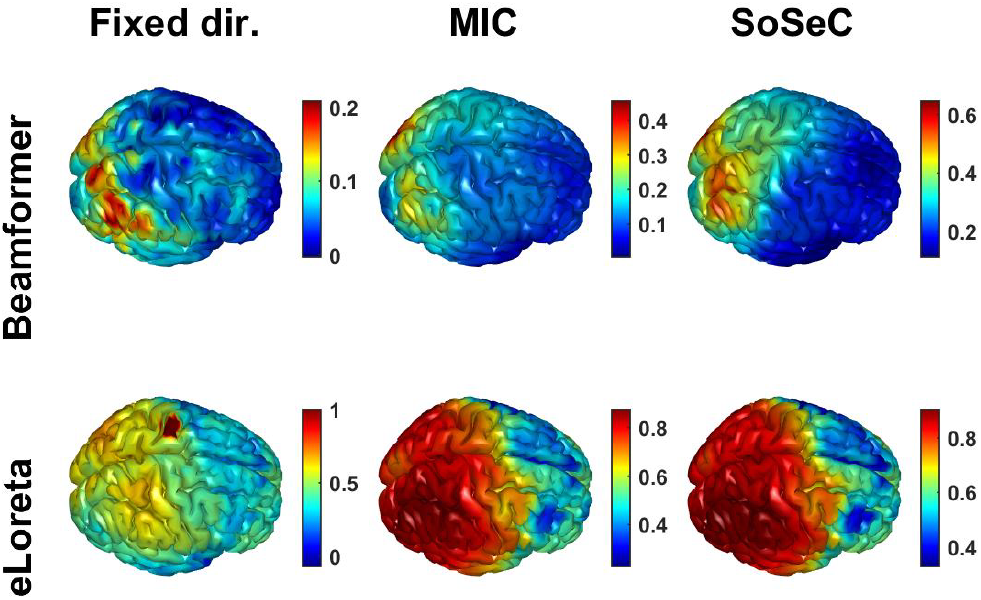
Results for three different coupling measures using two different inverse methods at 11 Hz calculated from the chosen example EEG data set. We observe interacting sources in occipital and parietal areas.

**Figure 7:**
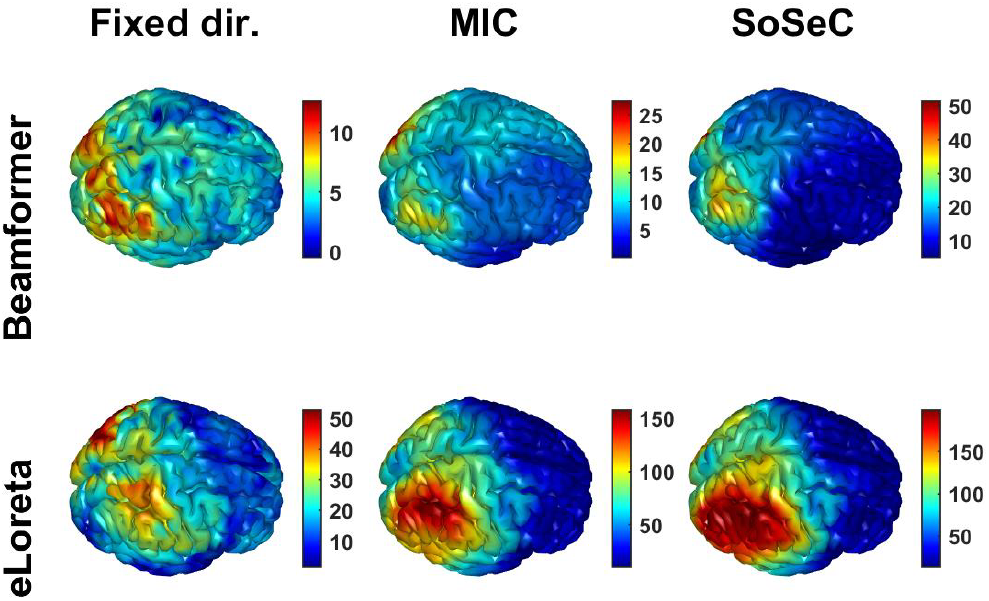
Corresponding z-scores for the coupling measures shown in Fig.6.

**Figure 8:**
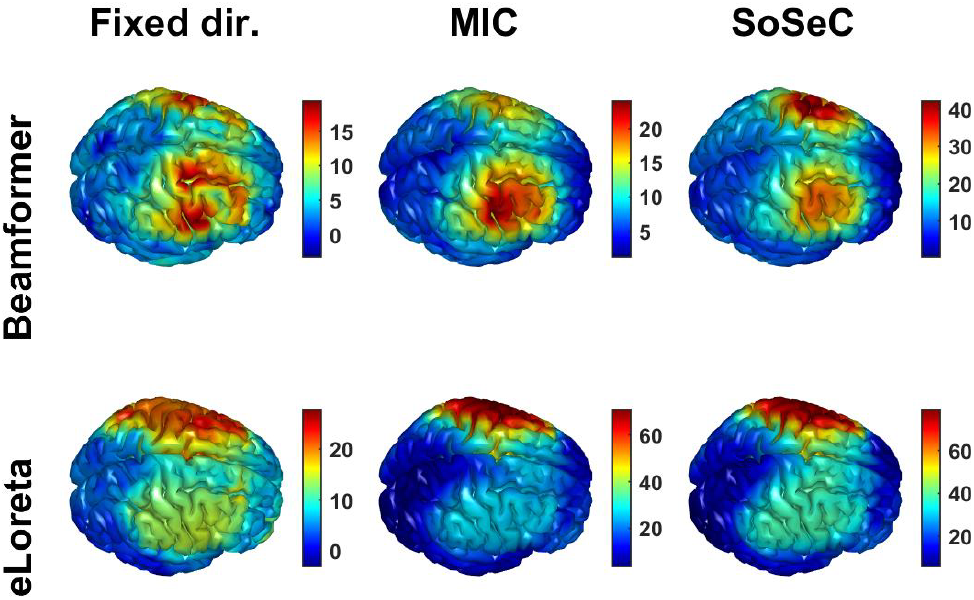
z-scores for three different coupling measures using two different inverse methods at 14 Hz calculated from the chosen example data set. We observe interacting sources in left and right motor areas.

### 3.3. EEG resting state data

To further characterize our new method, we applied it to resting-state EEG recordings (5-10min, sampled at 1 kHz, eyes closed) of 22 patients diagnosed with first-episode schizophrenia and 24 healthy controls. These data were provided by the Department of Psychiatry at the University Medical Center Hamburg-Eppendorf. EEGs were recorded using 64 Ag/AgCl electrodes positioned according to the 10–20 system and sampled at 1 KHz. Offline, artifacts were removed using independent component analysis (ICA). In addition, the data were down-sampled to 256 Hz and re-referenced to common average. For further details of the electrode positioning and preprocessing of the data we refer the reader to [Andreou et al., 2015a] and [Andreou et al., 2015b].

The preprocessed resting state data sets were then cut into epochs of 2 seconds and, using a segment length of 1 second and a segment shift of 0.5 seconds, Fourier-transformed after being tapered using a Hanning window. Data were analyzed in a frequency range from 1 to 100 Hz with a step size of 1 Hz. Based on the Fourier coefficients, we calculated cross spectra of the data and projected them to cortical source space using [Van Veen et al., 1997] as well as eLORETA (exact low-resolution brain electromagnetic tomography; [Pascual-Marqui, 2007, Pascual-Marqui et al., 2011]) filters. All computations were realized in a realistic 3-shell head model based on the MNI152 template brain (Montreal Neurological Institute; http://www.mni.mcgill.ca). Source connectivity was estimated within a continuous grid of 624 voxels and leadfields were calculated as described in [Nolte and Dassios, 2005]. For each voxel, we derived the maximum interaction with all other voxels (sensors for our new method, respectively) using three different coupling measures: (1) MIC (maximized imaginary coherence; Ewald et al., 2012), (2) a measure that fixes source orientations such that power is maximized which we label Fixed dir./Uni(var), and, (3) our new method SoSeC. All presented coupling measures show the imaginary part of coherency and are displayed in their squared form. Using a jackknife procedure, we also derived debiased versions of our coupling estimates (for details see above). In the following, we either display the debiased estimates of the three connectivity measures investigated or z values as derived by dividing coupling values by their respective standard deviations. Fig. 9 shows grand averages of our connectivity findings as a function of frequency. Whole-brain resting state coupling peaks in the alpha frequency range (8-12 Hz), for eLORTEA (Fig. 9 A) as well as beamformer (Fig. 9 B) filters, and results are most robust (highest z scores) for our new method SoSeC.

**Figure 9:**
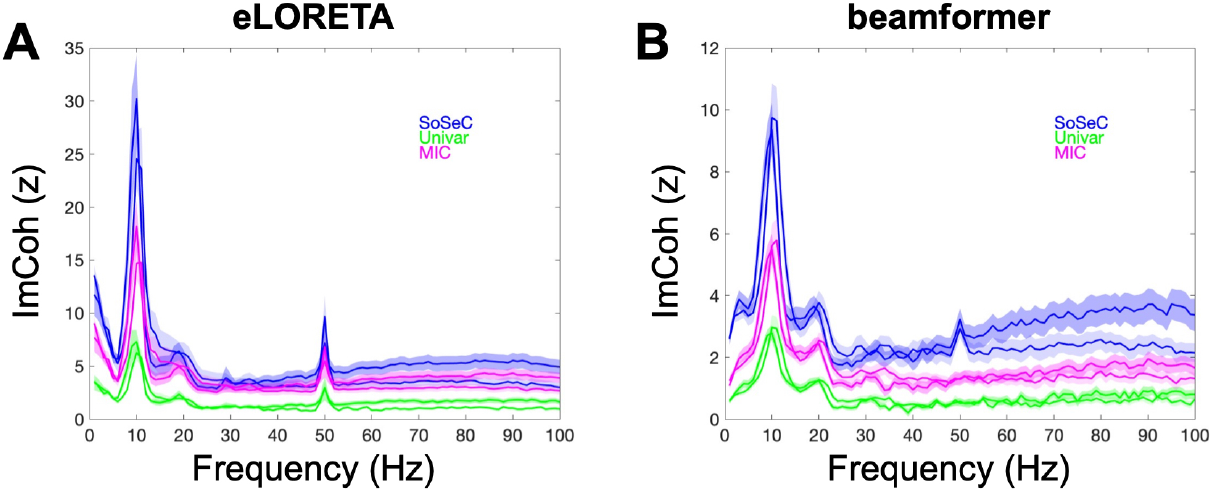
Grand average whole brain resting-state connectivity as a function of frequency (shown in Hz). Three different debiased coupling measures are shown: MIC (maximized imaginary coherence; Ewald et al., 2012), displayed in magenta, Univar, a measure that fixes source orientations such that power is maximized (see main text for details) plotted in green, and our new measure SoSeC (source-to-sensor-coupling) in blue. All connectivity estimates are shown in z values (imaginary coherence (ImCoh) values divided by their standard deviations). In A, values calculated using eLORETA filters are shown, in B beamformer filters are used. Shaded areas denote standard errors of the mean (SEM), darker shadows correspond to data from patients, lighter shadows to data from healthy control participants. Data shown are averaged across 22 patient datasets and 24 control participant datasets, respectively, as well as across 624 voxels. Resting-state connectivity peaks in the alpha frequency range (8-12 Hz).

To assess the impact of the regularization on the calculation of our new measure SoSeC, we analyzed one EEG dataset with different parameter combinations. For both, the first and the second regularization parameter, we created logarithmically spaced vectors with 10 values ranging from 10^−8^ to 101 as well as the value of 0.001 (which is the default for both regularization parameters). Each of these 121 (11 times 11) parameters configurations were used for the calculation of SoSeC values and their corresponding z values. For illustration purposes, we only show a selection of connectivity estimates employing these parameter configurations and plot them as a function of frequency (Fig.10).

**Figure 10:**
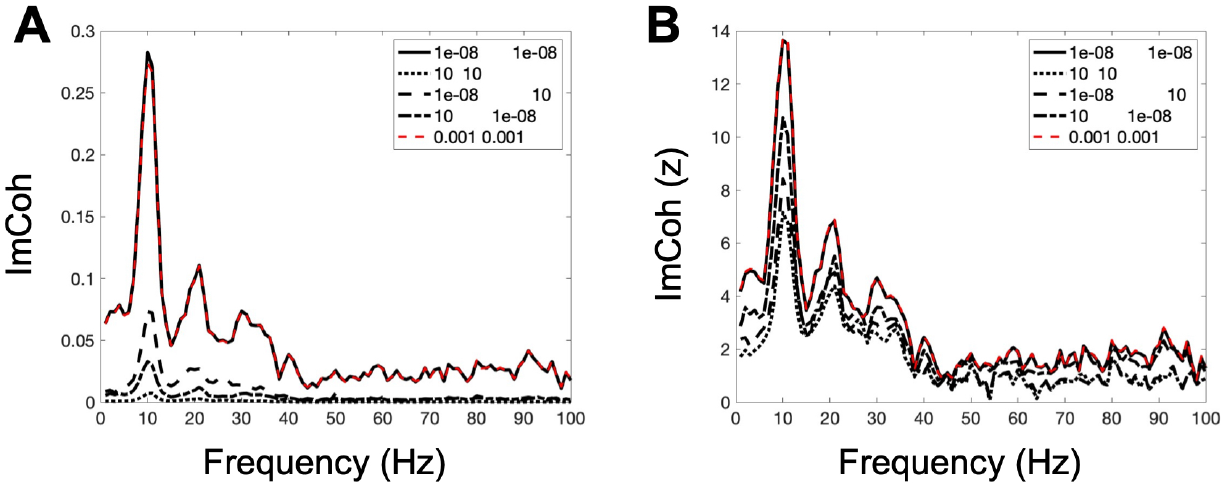
SoSeC (source-to-sensor-coupling) estimates as a function of frequency (shown in Hz) for different regularization parameter combinations. Connectivity estimates are displayed in imaginary coherence (ImCoh) raw values in A, and z values (ImCoh (z)) in B. Values were calculated for one EEG resting-state dataset of a single control participant and only beamformer filters were employed. Regularization parameter values of 0.001 (dashed red lines) appear to be a reasonable choice, whereas parameter values of 1 result in an almost zero connectivity pattern. Values shown are averages across 624 voxels.

To characterize the spatial distribution of the connectivity results, we exemplarily show restingstate coupling in the alpha band (8-12 Hz) for SoSeC, Univar, and MIC separately (Fig.11 A-C). Connectivity peaks in occipital cortex with visual areas being most strongly connected. In addition, we conducted repeated-measures correlation analyses on alpha coupling values, allowing for a statistical evaluation of the similarity between SoSeC and Univar on the one hand (Fig.11 D), and SoSeC and MIC on the other hand (Fig.11 E). The spatial patterns of the different connectivity measures investigated are very similar on a single patient dataset level, resulting in significant correlations. Again, z values were highest for our new method SoSeC. For the correlation analysis, we employed the software package rmcorr [Bakdash and Marusich, 2017], implemented in RStudio (RStudio, Inc., Boston, MA, 2020). For the rest of the analysis, custom-made Matlab (MathWorks, Natick, MA) code and functions implemented in our own toolbox METH were used.

**Figure 11:**
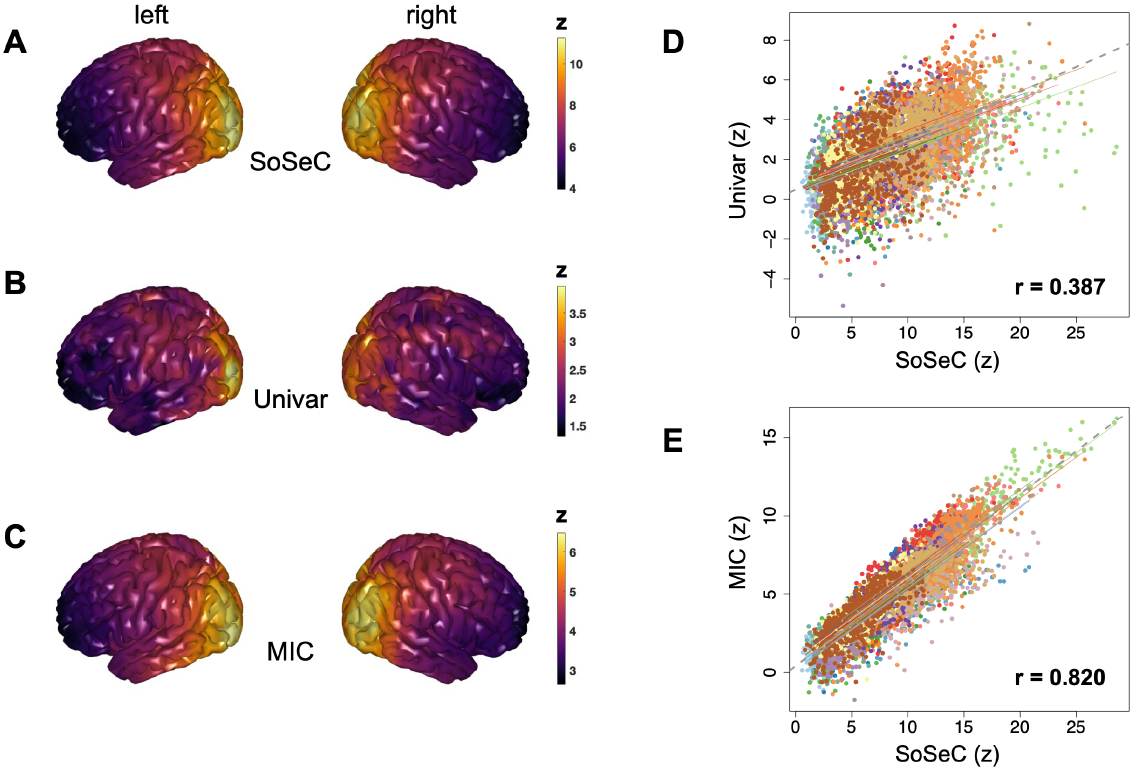
Source connectivity in the alpha frequency range (8-12 Hz). Three different debiased coupling measures are shown: MIC (maximized imaginary coherence; Ewald et al., 2012), displayed in A, Univar, a measure that fixes source orientations such that power is maximized (see main text for details) plotted in B, and our new measure SoSeC (source-to-sensor-coupling) in C. All connectivity estimates are shown in z values (imaginary coherence (ImCoh) values divided by their standard deviations). Only patient data are shown (average across 22 datasets), projected to the cortical surface. Warmer (yellow) colors indicate higher values. In D and E, repeated-measures correlations of different source connectivity estimates in the alpha frequency range (8-12 Hz) are presented. In D, the relation of SoSeC and MIC values (maximized imaginary coherence; Ewald et al., 2012; y-axis) is plotted. In E, data for SoSeC (x-axis) and Univar (y-axis) estimates are displayed. For both plots, z values are shown. Dot clouds of different colors show data of single patient datasets (n=22) for 624 voxels, complemented by their regression lines. The dashed line illustrates the correlation at the group level. Both correlations are statistically significant (*p <* 0.001), indicating similar spatial distributions for the different connectivity measures investigated.

## 4. Conclusion

We here propose a new method to detect interacting neuronal sources which is a) robust to artifacts of volume conduction, b) computationally efficient, and c) statistically robust. The basic procedure is to estimate the interaction of the activity in each source location with the signal in sensor space rather than with all other possible source locations. This dramatically reduces the computational cost. Since we only detect phase shifted interactions by calculating the imaginary part of coherency between source and sensor space, we can guarantee that this interaction does not reflect the ‘coupling’ of a signal with itself. The idea behind this approach is the trivial observation that any interaction which is observable between sources must also be contained in the sensors from which the corresponding source activities were calculated.

The result of this approach contains limited information on source interactions. We have an estimate for each source location whether the corresponding source is interacting, but we do not know with which other source or sources it is acting. This is similar to displaying a degree-map, i.e. for each voxel one displays the number of other voxels it interacts with [Van Vliet et al., 2018]. While conceptually the number of interactions is in general very different from the strength, we expect those differences to be small in practice because inverse solutions are always blurred and strong interactions will result in many estimated interactions. In practice, the information about coupling between all possible pairs of sources is useless for a large number of grid points, if this information is not condensed to a tractable form. E.g., a typical full brain grid with 1cm spacing contains around 5000 voxels, and the coupling between all pairs consists of 25 million numbers, which are, in addition, impossible to visualize because each number represents a point in a 6-dimensional space corresponding to two locations in a 3-dimensional space. To visualize the full information one can e.g. maximize the coupling for each seed with respect to all other sources. This reduction of information eventually leads to the analogous information as we get with the proposed approach SoSeC directly, with a computational cost being typically hundreds of times smaller than by first calculating coupling between all pairs using a multivariate optimization approach [Ewald et al., 2012].

We emphasize that results from SoSeC are similar but not identical to classical approaches. While for classical approaches one assumes that both sources a dipolar, this is not the case for SoSeC where only one of the sources is assumed to be dipolar. However, this distinction might be rather academic as neuronal sources are typically well described by dipoles. In practice, statistical robustness is the more relevant question. We found that SoSeC is statistically more robust, i.e. z-scores were higher, both in comparison to a classical univariate approach (i.e. one time series for each source location) and to a classical multivariate approach (i.e. 3 time series for each source direction). The computational advantage of SoSeC over classical methods only applies to multivariate approaches. However, we also found that the univariate approach is far less robust than the multivariate approach we tested here which therefore cannot be recommended.

Of course, SoSeC can be considered as a first step in a more in-depth analysis: one or more local maxima of a SoSeC result can be considered as seeds for a classical approach. This may then result in all relevant information for pairs of coupling while preserving the low computational cost for the whole analysis. Another modification of the analysis pipeline could be to first find a local maximum and then to project out the corresponding pattern in the data in signal space and restart SoSeC on that. The result is the coupling of sources to anything except this seed. We here only present a few ideas on further developments, which are, however, beyond the scope of this paper.

## Acknowledgment

This research was partially funded by the BMBF (161A130), the German Research Foundation (DFG, SFB936/A2/A3/Z3 and TRR169/B1/B4 and SPP2041/EN533/15-1), and from the Landesforschungsförderung Hamburg (CROSS, FV25). We also acknowledge support by the European Union (project cICMs, ERC-2022-AdG-101097402). Views and opinions expressed in this paper are those of the authors only and do not necessarily reflect those of the European Union or the European Research Council. Neither the European Union nor the granting authority can be held responsible for them.

